# Maintenance of fertility in the face of meiotic drive

**DOI:** 10.1101/675108

**Authors:** Lara Meade, Sam Finnegan, Ridhima Kad, Kevin Fowler, Andrew Pomiankowski

## Abstract

Selfish genetic elements that gain a transmission advantage through the destruction of sperm have grave implications for drive male fertility. In the X-linked SR meiotic drive system of a stalk-eyed fly, we found that drive males have greatly enlarged testes and maintain high fertility despite the destruction of half their sperm, even when challenged with fertilising large numbers of females. Conversely, we observed reduced allocation of resources to the accessory glands that probably explains the lower mating frequency of SR males. Body size and eyespan were also reduced, which are likely to impair viability and pre-copulatory success. We discuss the potential evolutionary causes of these differences between drive and standard males.

## Introduction

Meiotic drive genes gain a transmission advantage through manipulation of meiosis or gametogenesis and are likely to have profound ecological and evolutionary consequences, ranging from the evolution of sex determination systems and changes in karyotype, to impacts on population persistence and sexual selection (Hurst and Werren 2001; Jaenike 2001; Werren 2011; Lindholm et al. 2016). Drivers have been uncovered in a wide range of taxa, with a preponderance of linkage to the sex chromosomes in the heterogametic sex (Hurst and Pomiankowski 1991; Jaenike 2001; Taylor and Ingvarsson 2003). When meiotic drive occurs in males, it severely disrupts the maturation and fertilisation capacity of non-carrier sperm, imposing a fertility disadvantage to organismal fitness (Price and Wedell 2008) which is exaggerated under conditions of sperm competition (Taylor et al. 1999; Angelard et al. 2008; Price et al. 2008*a*) and typically has pleiotropic viability costs in both sexes (Burt and Trivers 2006).

The extent to which these and other detrimental effects of sperm-killer drive promote adaptive responses in the host species has received limited attention. There is an extensive literature on genetic elements that interfere and suppress the action of drive. For example, in *Drosophila* species, suppressors of X-linked drive have been found on the Y chromosome (Carvalho et al. 1997; Cazemajor et al. 1997; Branco et al. 2013) and throughout the rest of the genome (Carvalho and Klaczko 1993; Atlan et al. 2003; Tao et al. 2007). A more recent suggestion is that drive may promote the evolution of female polyandry in order to dilute the ejaculates of drive males (Haig and Bergstrom 1995; Zeh and Zeh 1997; Wedell 2013). There is some evidence for this from experimental evolution studies using populations exposed to meiotic drive in *D. pseudoobscura* (Price et al. 2008*b*) and *Mus musculus* (Manser et al. 2017), and from natural populations in which the rate of multiple mating correlates negatively with the frequency of drive in *D. pseudoobscura* (Price et al. 2014) and *D. neotestacea* (Pinzone and Dyer 2013). Female mate choice may additionally evolve in response to drive. In stalk-eyed flies, meiotic drive has been linked to small eyespan, which may allow females to avoid mating with carrier males through assessing eyespan (Wilkinson et al. 1998*b*; Cotton et al. 2014). Female house mice could avoid mating with drive males through detecting unique major histocompatibility alleles linked to the driving *t* complex (Silver 1985; Lindholm et al. 2013), although evidence remains unclear (Lindholm and Price 2016).

Another, as yet unexplored, route by which males could adapt to drive is by increasing the allocation of resources to sperm production, to offset the destructive effect of drive on gametogenesis. Sperm number is positively correlated with testis size in many intra-specific studies (Gage 1994; Fry 2006; Hettyey and Roberts 2006) and increased testis size is a well characterised evolutionary response to heightened sperm competition favouring greater sperm production (Hosken and Ward 2001; Pitnick et al. 2001; Simmons and García-González 2008; Gay et al. 2009). The loss of sperm in drive males could be compensated for by increased investment in testis. Meiotic drive elements are typically found within inversions or other areas of low recombination that keep drive and insensitive responder loci together (Palopoli and Wu 1996; Johns et al. 2005; Dyer et al. 2007), facilitate the spread of modifiers that enhance transmission distortion (Hartl 1975; Larracuente and Presgraves 2012) and are predicted to be enriched for male beneficial sexually-antagonistic alleles (Rydzewski et al. 2016). For similar reasons, alleles that enable compensatory investment in testes could become associated with the drive haplotype.

We test this idea using the Malaysian stalk-eyed fly species *Teleopsis dalmanni*. This species harbours SR, an X-linked driver, which produces strongly female-biased broods due to the destruction of Y-bearing sperm (Presgraves et al. 1997; Wilkinson and Sanchez 2001). Meiotic drive arose around 2-3.5 Mya in the *Teleopsis* clade, and the X^SR^ drive chromosome in *T. dalmanni* is estimated to have diverged from a non-driving ancestor (X^ST^) around 1 Mya (Swallow et al. 2005; Paczolt et al. 2017), and is characterised by a large inversion(s) covering most of the X chromosome (Johns et al. 2005; Paczolt et al. 2017). X^SR^ is found at appreciable frequencies (10 – 30%) across populations and generations (Wilkinson et al. 2003; Cotton et al. 2014) but appears to lack genetic suppressors (Reinhold et al. 1999; Wolfenbarger and Wilkinson 2001; Paczolt et al. 2017). This means that there has been ample time and opportunity for adaptive responses to selection to evolve in male carriers of the drive chromosome.

We determined whether SR and standard (ST) males differed in their reproductive (testis and accessory gland size) and morphological traits (eyespan and body size). Testis size predicts the amount of sperm found within female storage (Fry 2006). Accessory glands produce all non-sperm components of the ejaculate, and accessory gland size is positively associated with male mating frequency (Baker et al. 2003; Rogers et al. 2005*a*, 2005*b*). Body size and eyespan are also important predictors of male mating frequency (Wilkinson et al. 1998*a*; Small et al. 2009; Cotton et al. 2010). We determined SR and ST sperm production by mating them to low or high numbers of females over a 10-hour period, counting the number of fertilized eggs produced. Males were also exposed to females over a short time period (30-minutes) to compare the copulation rate of SR and ST males.

## Methods

Details of stock collection and day-to-day upkeep can be found in the Appendix. Experimental males were taken from the SR-stock population in which males are a ~50:50 mix of X^SR^ and X^ST^ genotypes. Experimental females were taken from the ST-stock population, which lacks meiotic drive. Single non-virgin males were allowed to mate freely with either one or five virgin ST-stock females, over a period of 10 hours. Mated females were allowed to lay eggs for 14 days, by which time most females had stopped laying fertile eggs. Fecundity was recorded through egg counts, and egg hatch was used as an estimate for fertility. On the following day, experimental males, and a similar number of unmated males, were anaesthetised on ice and their testes and accessory glands were removed (fig. 1A) and photographed under differential interference contrast microscopy. Organ area was measured at x50 magnification by tracing the outline. Male eyespan (Hingle et al. 2001) and a proxy for body size, thorax length, (Rogers et al. 2008) were measured.

**Figure 1.**
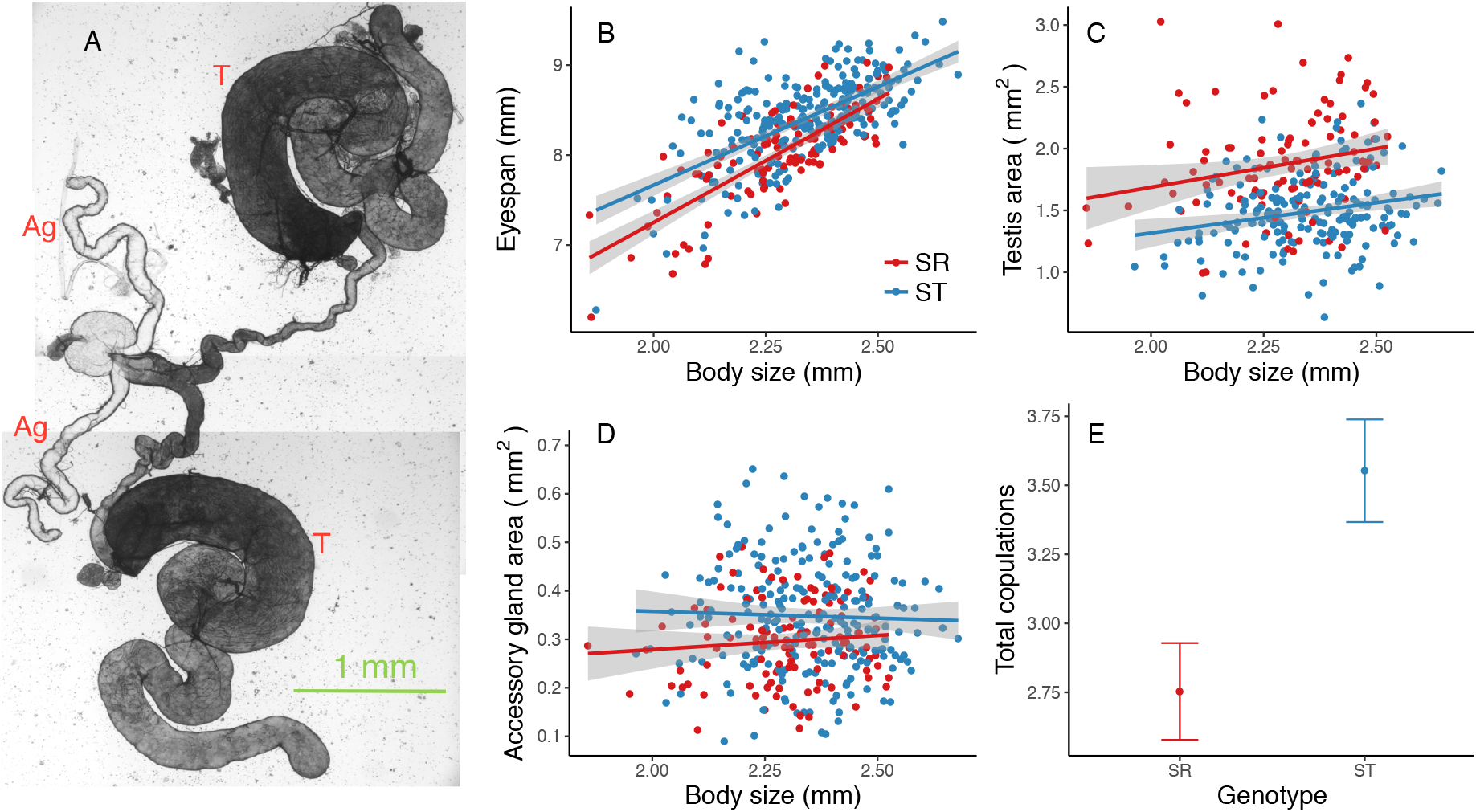
*A*, testes (T) and accessory glands (Ag) after dissection. *B–D*, male morphological and reproductive trait size for SR (red) and ST (blue) males, plotted against male body size. *B*, male eyespan. *C*, male testis area. *D*, male accessory gland area. SR males have smaller body size, eyespan and accessory gland size, but larger testis size. Grey shading shows ± s.e. *E*, mating frequency, measured as total number of copulations (mean ± s.e.) observed over 30 minutes.

In a second experiment, SR-stock males were introduced to two ST-stock non-virgin females at artificial dawn. All copulations were counted during 30 minutes. To minimise any effects on mating frequency due to female choice, the experimental males were standardised to have a narrow range of eyespan (7.5 – 8.5 mm).

Males from both experiments were genotyped using either two X-linked INDEL markers, *comp162710* and *cnv395*, or a microsatellite marker, *ms395*. Allele size of these markers reliably indicates the SR genotype of the males in our laboratory stocks (Meade et al. 2018).

### Statistical analysis

We tested if male genotypes differed in their morphological (body size and eyespan; linear models) and reproductive traits (testis size and accessory gland size; linear mixed effects models). Differences in relative trait sizes between genotypes, as well as in absolute trait sizes (models where body size is excluded) are reported. The total number of fertile eggs (Poisson generalised linear mixed effects model (GLMM)) and proportion fertility (fertile eggs, non-fertile eggs; binomial GLMM) of females are compared when mated to SR (i.e. XSR/Y genotype) or ST (i.e. XST/Y genotype) males. We also tested if male reproductive traits, and their interaction with male genotype, were important predictors of fertility. Lastly, we tested whether SR and ST males differed in their mating frequency over 30-minutes by comparing the likelihood that SR and ST males mate at all (binomial GLMM), as well as the total number of copulations among males that mated at least once (Poisson GLMM).

To avoid collinearity of male morphological and reproductive traits with body size, models used residual values (Dormann et al. 2013). Where appropriate, experimental batch was included as a random effect. Further details and model effect sizes can be found in the Appendix.

## Results

### SR trait size

SR males had small body size (mean ± s.e. 2.290 ± 0.013 mm) compared to ST males (2.336 ± 0.009 mm; F_1,357_ = 8.745, P = 0.003; fig. 1B). SR males also had small absolute (SR: 8.048 ± 0.046mm; ST: 8.402 ± 0.031mm; F_1,357_ = 42.631, P < 0.001; fig. 1B) and relative eyespan (F_1,355_ = 0.713, P = 0.016), especially when body size was small (body size by genotype interaction F_1,355_ = 4.175, P = 0.042).

Despite their small body size, SR testis size was large (1.940 ± 0.050 mm^2^) compared to ST males (1.54 ± 0.028 mm^2^; F_1,280.16_ = 73.796, P < 0.001; fig. 1C). SR males also had large relative testis size (F_1,282.78_ = 99.982, P < 0.001). In contrast, SR males had small absolute (SR: 0.306 ± 0.011 mm^2^; ST: 0.348 ± 0.010 mm^2^; F_1,335.36_ = 16.353, P < 0.001; fig. 1D) and relative accessory gland size (F_1,334.03_ = 7.801, P = 0.006). Taking relative values for each genotype, eyespan (F_1,286_ = 19.892, P <0.001) and accessory gland size (F_1,274.418_ = 26.008, P <0.001) increased with testes size, but the rate was reduced in SR males (interaction eyespan: F_1,286_ = 5.261, P = 0.023, fig. A1; interaction accessory glands: F_1,268_ = 8.375, P = 0.004, fig. A2).

### SR fertility

SR males did not differ from ST males in total (mean ± s.e. SR: 112.047 ± 8.290, ST: 107.053 ± 5.597; χ^2^_1_ = 2.416, P = 0.120, N = 215; fig. 2A, 2B) or proportion fertility (SR: 0.833 ± 0.025, ST: 0.762 ± 0.019; χ^2^_1_ = 2.469, P = 0.116, N = 215) when kept with females over an extended 10-hour period. Males mating with five females achieved higher total fertility (one female: 79.231 ± 5.090, five females: 138.123 ± 6.653; χ^2^_1_ = 43.698, P < 0.001, N = 215) but a lower proportion fertility (one female: 0.804 ± 0.024, five females: 0.763 ± 0.199; χ^2^_1_ = 6.021, P = 0.014, N = 215) than those mating with a single female. The interaction between mating group (one or five females) and genotype did not influence total (χ^2^_1_ = 0.591, P = 0.442, N = 215) or proportion fertility (χ^2^_1_ = 1.377, P = 0.241, N = 215).

**Figure 2.**
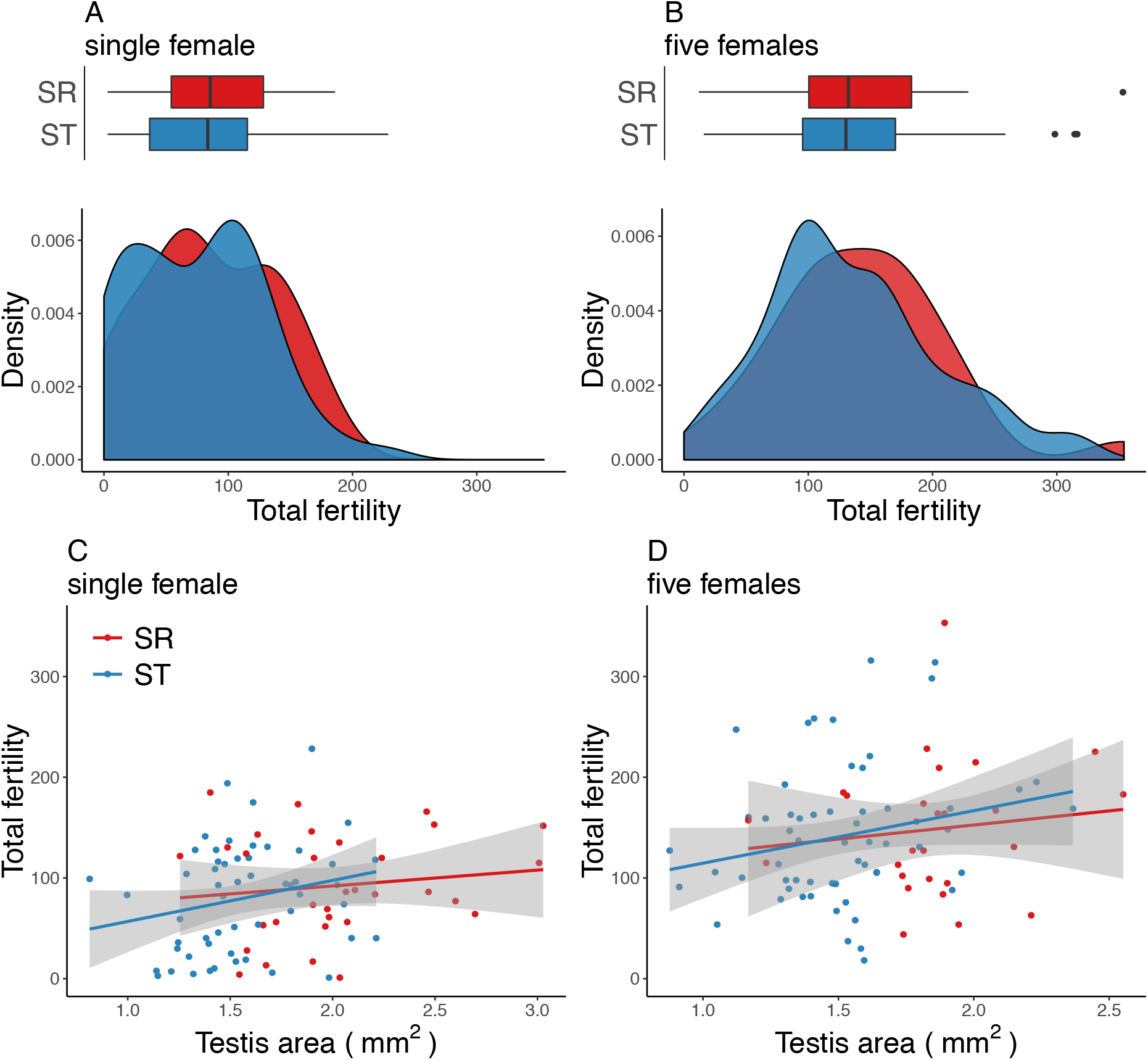
*A–B, upper:* box plots (median and interquartile range) and *lower:* Kernel probability density of measures of total fertility of SR (red) and ST (blue) males. *A*, mated to a single female. *B*, mated to five females. Across both mating regimes, SR and ST males did not differ in the number of eggs fertilised. *C–D*, absolute testis area plotted against total fertility. *C*, mated to a single female. *D*, mated to five females. Across both mating regimes, total fertility increased with testis area in SR and ST males. Grey shading shows ± s.e.

Male testis size was an important predictor of fertility. Both total (χ^2^_1_ = 5.897, P = 0.015, N = 165; fig. 2C, 2D) and proportion fertility (χ^2^_1_ = 18.837, P < 0.001, N = 165) were greater amongst males with larger testis size, even when accounting for male body size (total: χ^2^_1_ = 6.216, P = 0.013, N = 165; proportion: χ^2^_1_ = 16.646, P < 0.001, N = 165). The addition of testis size did not alter the relationship between genotype and total (χ^2^_1_ = 0.018, P = 0.895, N = 173) or proportion fertility (χ^2^_1_ = 0.260, P = 0.610, N = 173). There was no interaction between testis size and genotype predicting total (χ^2^_1_ = 0.164, P = 0.686, N = 173) or proportion fertility (χ^2^_1_ = 0.617, P = 0.432, N = 173). Accessory gland size did not predict total (χ^2^_1_ = 0.032, P = 0.858, N = 165) or proportion fertility (χ^2^_1_ = 0.160, P = 0.689, N = 165).

### SR mating frequency

A total of 493 copulations from 193 males were observed over the 30-minute mating trials. SR males (mean ± s.e. 2.750 ± 0.175, N = 81) copulated fewer times on average than ST males (3.550 ± 0.186, N = 76; χ^2^_1_ = 6.304, P = 0.012; fig. 1E), but were not less likely to mate at least once (SR: 81/104, ST: 76/89; χ^2^_1_ = 1.665, P = 0.197, N = 193).

## Discussion

One of the main features of drive in males is reduced sperm production due to the dysfunction of non-carrier sperm. This has been reported to cause a loss in fertility in a variety of species including *Drosophila* (Hartl et al. 1967; Jaenike 1996; Angelard et al. 2008; Price et al. 2012; Pinzone and Dyer 2013), house mice (Carroll et al. 2004), and *Silene alba* (Taylor et al. 1999). Here, we present evidence that SR males in *T. dalmanni* overcome this deficit by having greatly enlarged testes. SR males carry an extreme form of the X^SR^ drive chromosome, siring female-only broods due to the dysfunction of Y-bearing gametes. Despite gamete loss, SR males achieve fertility at a level equivalent to that of ST males, both when exposed to a single female or 5 females over a 10-hour period (fig. 2). Our results contradict a previous study which found an SR fertility deficit using a similar design (Wilkinson et al. 2006). But this study measured fertility as the number of adults that eclosed, compounding fertility with egg-to-adult survival. Recent work shows larval survival is reduced in drive heterozygous females (Finnegan et al. 2019), which could account for the drop in SR male fertility. The patterns in *T. dalmanni* are in contrast to other insect species with X-linked meiotic drive which generally show a deficiency in fertility of drive males either after a single or multiple matings (Jaenike 1996; Atlan et al. 2004; Angelard et al. 2008; Price et al. 2012; Pinzone and Dyer 2013).

These experiments were designed to test whether daily sperm reserves differ between SR and ST males, not to replicate normal levels of mating observed under natural conditions which occur at far lower rates (Cotton et al. 2015). On dissection, we discovered that SR males have greatly enlarged testes (fig. 1C), about 26% larger than ST males. This difference remained after controlling for body size (fig. 1C). Our interpretation is that the increase in testis size allowed SR males to compensate for the loss of sperm due to the action of meiotic drive. This is supported by the finding that fertility increased with increasing testis size, both for absolute and relative testis size, in both SR and ST males (fig. 2). Our interpretation also aligns with previous findings that SR male ejaculates deliver similar numbers of sperm as ST males, after single and multiple matings (Meade et al. 2018). Despite the destruction of half their sperm, the increased investment in SR testis size (i.e. sperm production) allows them to deliver sufficient sperm to achieve similar fertility as ST males. To further understand the extent of this compensation, we need to assess SR male success under sperm competition, which is the norm in *T. dalmanni* (Wilkinson et al. 1998*a*; Baker et al. 2001; Corley et al. 2006). Previous work suggests that SR males perform poorly under sperm competition (Wilkinson et al. 2006) but this assessment again does not take account of the lower egg-to-adult viability of X^SR^ carriers (Finnegan et al. 2019) which could simulate an advantage of ST males in sperm competition. In our experimental design, autosomal background was standardised across SR and ST males. So, it seems likely that control of testis size is linked to alleles that are located in the X^SR^ chromosomal inversion and that such alleles arose as an adaptive response to sperm dysfunction caused by drive, but further investigation is needed to establish this view.

We found morphological trait divergence in accessory gland size, which are small in SR males, even after controlling for body size (fig. 1D). Previous work in *T. dalmanni* shows that accessory gland size is linked with the mating rate (Baker et al. 2003; Rogers et al. 2005*a*).

This might explain why the mating frequency of SR males was low, being about 75% of the rate for ST males over a 30-minute period (fig. 1E). In addition, SR males have small body size and small eyespan for their body size (fig. 1), traits likely to reduce male mating success, both in male-male agonistic interactions (Panhuis and Wilkinson 1999; Small et al. 2009) and in attracting and mating with females (Wilkinson and Reillo 1994; Hingle et al. 2001; Cotton et al. 2010). The increased allocation of resources to testes in SR males potentially causes a reduction in the resources available for investment in accessory glands, as both traits develop over several weeks post-eclosion (Baker et al. 2003; Rogers et al. 2008). Resource competition with testes is not an obvious reason for reduced body size and eyespan which are determined during larval development. However, expression of these traits might be connected via juvenile hormone which has been shown to mediate a trade-off between eyespan and testes in stalk-eyed flies (Fry 2006).

Small body size and eyespan are also likely to arise from the low genetic condition of drive males. The *T. dalmanni* SR inversion(s) covers nearly all of the X chromosome, capturing one third of the stalk-eyed fly genome (Johns et al. 2005; Paczolt et al. 2017). X^SR^ alleles will be subject to weak natural selection due to reduced recombination and liable to accumulate deleterious mutational effects (Kirkpatrick 2010). Consistent with a lack of recombination, there are 955 fixed sequence differences between transcripts linked to X^SR^ and X^ST^ (Reinhardt et al. 2014). Such mutations are expected to have a negative effect on costly, condition-dependent traits, such as body size and eyespan, whose expression is affected by multiple loci distributed throughout the genome (David et al. 2000; Cotton et al. 2004; Bellamy et al. 2013). Given SR males have small eyespan, they will be unattractive and gain fewer mating opportunities. Consequently, investment in accessory glands which enable higher mating rates will give lower returns than the diversion of resources into larger testes which allow SR males to produce ejaculates of equivalent size to those of ST males, and be able to compete under the conditions of high sperm competition seen in stalk-eyed flies. These ideas about linking resource allocation, condition and mating rates need further investigation, in particular under the mating conditions that occur in the wild.

Here we demonstrate for the first time that through investment in testis, drive males can maintain fertility, despite sperm destruction. Other responses to drive, such as genetic suppression, polyandry and female choice, reduce the transmission advantage gained by drive, and lead to reductions in the equilibrium frequency of drive (Hartl 1975; Taylor and Jaenike 2002; Holman et al. 2015). In sharp contrast, increased investment in sperm production intensifies the transmission of drive, because the fertility gain to the individual male is also beneficial to the drive element itself. Such an association with meiotic drive has neither been theoretically modelled nor empirically studied previously, but has implications for the spread and equilibrium frequency of drive in natural populations.

## Authors’ contributions

All authors contributed to conceiving the project and methodology; LM and RK collected data on fertility and morphology; SF collected data on mating frequency; LM analysed the data; LM, KF and AP led the writing of the manuscript. All authors contributed critically to the drafts and gave final approval for publication.

## Acknowledgements

Funding was provided by a NERC Studentship held by LM, and by EPSRC (EP/F500351/1, EP/I017909/1) awards to AP and NERC grants (NE/G00563X/1, NE/R010579/1) to KF and AP.

## Data accessibility

Data will be uploaded to the Dryad Digital Repository

## Ethical statement

No ethical approval was required for this research

