## Appendix for "Maintenance of fertility in the face of meiotic drive"

### Contents

**1 Stock source and maintenance** **3**

**2 SR fertility and reproductive organ size** **4**

**3 SR mating frequency** **6**

**4 Genotyping** **6**

**5 Statistical analysis** **7**

**6 Model tables and effect sizes** **10**

**7 Figures** **29**

### 1 Stock source and maintenance

The stock populations were kept at 25°C, with a 12:12 h dark:light cycle and fed puréed sweetcorn twice weekly. Fifteen-minute artificial dawn and dusk periods were created by illumination from a single 60-W bulb at the start and end of the light phase. Experimental flies were collected from egg-lays placed in stock population cages. Egg-lays consist of damp cotton-wool and excess puréed sweetcorn contained in a Petri dish. After eclosion, adult flies were measured for eyespan and thorax length using ImageJ (v1.46) and separated by sex prior to sexual maturity (<3 weeks after eclosion). Upon entering an experiment individuals were all >6 weeks old and sexually mature (Baker et al. 2003). Experimental males were sourced from the sex-ratio meiotic drive stock population (SR-stock) and were housed in 12L mixed-sex cages with a 1:1 ratio males:females for at least 12 days.

##### *Standard stock population*

The standard stock (ST-stock) population was created from collections in 2005 (by S. Cotton and A. Pomiankowski) from the Ulu Gombak valley, Peninsular Malaysia (3°19'N 101°45'E) and maintained in high density cages (> 200 individuals) to minimise inbreeding. This population has been regularly monitored and does not contain meiotic drive and carries non-distorting standard X chromosomes ($X^{ST}$).

##### *Sex-ratio meiotic drive stock population*

Flies were collected in 2012 (by A. Cotton and S. Cotton) from the Ulu Gombak valley to create a sex ratio meiotic drive stock (SR-stock) population. To establish and maintain a stock with meiotic drive, a standard protocol was followed (Presgraves et al. 1997).

Briefly: wild males (of unknown genotype) were mated to ST-stock females and their offspring (F1) were collected. When an F1 brood was female biased (80% female, > 10 offspring), it was assumed that the father was a carrier of the sex-ratio distorting $X^{SR}$ chromosome, so that the F1 female offspring had genotype $X^{SR}/X^{ST}$. These females were mated with ST-stock males and their offspring (F2) were collected. Male F2 offspring are expected to be 50:50 $X^{SR}/Y$:$X^{ST}/Y$ as they inherit either an $X^{SR}$ or $X^{ST}$ chromosome from their mother. F2 males were subsequently mated to ST-stock females to identify those males carrying $X^{SR}$, and the process repeated.

Even though there was error in the assignment of individuals as carriers of $X^{SR}$, the process maintains the $X^{SR}$ chromosome in this stock. Over generations the SR phenotype has become more distinct as the stock maintenance procedure selects for female biased broods, so most biased offspring sex-ratios are now 100% female, or at least > 95% female biased (Meade et al. 2018). Note that because the SR-stock maintenance involves back-crossing to ST-stock males and females, the autosomes, Y-chromosome and mitochondrial backgrounds are homogenised across the two stocks. We hereafter refer to $X^{SR}/Y$ and $X^{ST}/Y$ males as SR and ST males respectively.

### 2 SR fertility and reproductive organ size

Non-virgin experimental males were housed for two days in single-sex cages. Males were all 7–8 weeks old. On the evening of the final day, two thirds of males were transferred to mating pots (Cotton et al. 2015). In the mating pots, males were housed individually in the upper chamber. In the lower chamber, separated from the male by a card partition, either one or five virgin females were added. These females were sourced from the ST-stock and had medium–large eyespan ($\geq$ 5.7mm). Males were otherwise moved individually to 500ml pots, without females. Just before artificial dawn the following day, card partitions were removed allowing interactions between males and females in the mating pots. After 10 hours, males and females were separated.

On the subsequent dawn, all males were anaesthetised on ice for dissection. Male testes and accessory glands were dissected into a small amount of PBS on a glass microscope slide and a cover slip added (Baker et al. 2003; Rogers et al. 2005; 2008). Uncoiled organs were visualised using differential interference contrast microscopy and images were photographed at x50 magnification using a monochrome microscope camera and QCapture Pro imaging software (v7.0). Organ area was measured using ImageJ (v1.50i) by tracing the outline of the organ to give a longitudinal surface area. The area of a single randomly chosen testis was measured and the area of both accessory glands were measured (Baker et al. 2003; Rogers et al. 2005). Males were stored in ethanol at -20°C prior to genotyping.

Immediately after the 10 hour mating period, females were transferred to 500ml pots and allowed to lay eggs. The bases of pots consisted of damp cotton-wool covered with blue tissue paper and females were fed with puréed sweetcorn. Eggs were collected every 2–3 days for 14 days, and were allowed to develop in a Petri dish with moist cotton-wool for at least a further 3 days. Eggs were observed under a light microscope at x10 magnification; fertilised eggs that have hatched appear as empty chorion cases. Additionally, eggs were counted as fertilised if they failed to hatch but showed clear signs of development (brown horizontal striations in the chorion and early mouth part formation, Baker et al. 2001). Unfertilised eggs show no signs of development.

### 3 SR mating frequency

Mating pots were set up in the afternoon before mating trials. Non-virgin experimental males were placed individually into the upper chamber and two non-virgin females in the the lower chamber, separated by a card partition. On the following artificial dawn, the partition was removed, allowing interactions between males and females. Males were observed for the following 30 minutes and the number of copulations were recorded. A successful copulation was defined as intromission lasting more than 30 seconds, as copulations shorter than this duration do not result in spermatophore transfer (Rogers et al. 2006). Males that attempted to mate but were unsuccessful (defined as following and attempted mounting, but no intromission) were presented with a different set of females and observed for an additional 30 minutes. Males were stored in ethanol at -20°C prior to genotyping.

### 4 Genotyping

Two INDEL markers, *comp162710* and *cnv395*, or a microsattelite marker, *ms395*, were used to identify SR and ST males. Allele sizes for these markers segregate into two size classes. The presence of these markers in males is a good indicator of whether they are carriers of the $X^{SR}$ or $X^{ST}$ chromosome and produce significantly biased brood sex-ratios (Meade et al. 2018).

A standard protocol was followed to extract DNA (Bruford et al. 1998). For each sample, half a thorax was crushed and digested in 250$\mu$l digestion solution (20mM EDTA, 120mM NaCl, 50mM Tris-HCL, 1% SDS, pH 8.0) and 10$\mu$l proteinase K (10mg ml$^{-1}$), and the samples incubated for $\sim$12hrs at 55°C. Proteins were precipitated out with 300$\mu$l of 4M ammonium acetate and spun at 13000rpm for 10min. The supernatant was aspirated into 1ml absolute ethanol to precipitate out the DNA, which was pelleted by spinning at 13000rpm for 10min. The DNA pellet was washed in 500ml of 70% ethanol and allowed to dry before being stored in 50$\mu$l T10 E0.1 buffer at -20°C. PCR reactions were performed on a 2720 Thermal Cycler (Applied Biosystems, Woolston, UK) in 2$\mu$l samples, containing 1$\mu$l QIAGEN Mastermix (QIAGEN, Manchester, UK), 1$\mu$l Primer mix and 1$\mu$l DNA (dried). All primers were at a 0.2$\mu$M concentration. PCR reactions had an initial denaturing stage of 95°C for 15min followed by 45 cycles of 94°C for 30sec, 60°C for 1min 30sec and 72°C for 1min 30sec. This was completed by an elongation step of 60°C for 30min. The Applied Biosystems ABI3730 Genetic Analyzer was used to visualise the microsatellites, with a ROX500 size standard. Allele sizes were assigned using GENEMAPPER 4.0, or in R (v3.32, R Core Team 2016) using the *Fragman* package (v1.07, Covarrubias-Pazaran et al. 2016). Sequencing work was carried out at the NERC Biomolecular Analysis Facility at the University of Sheffield.

### 5 Statistical analysis

To avoid collinearity of male morphological and reproductive traits with body size, models used residual values for eyespan, testis size and accessory gland size. For example, residual eyespan are the residuals from an linear model (LM) after the variation in eyespan explained by body size has been removed (Dormann et al. 2013). Reported P-values were calculated with type II tests using the *Anova* function from the *car* package, and specifying type III tests where significant interactions were present (Fox and Weisberg 2011).

#### Body size and eyespan

To test if male genotypes differed in body size (using thorax length as a proxy), body size was analysed as as a function of genotype in a LM. Body size was first transformed (squared) to normalise the distribution of the errors. Eyespan was modelled in an LM as a function of body size, genotype and their interaction.

#### Reproductive organ size

To test if male testis and accessory gland size differed between genotypes, we analysed both testis size and accessory gland size as functions of mating group (one female, five females or unmated), body size, eyespan and genotype (SR or ST), and up to their three-way interactions in linear mixed effect models (LMMs), including batch as a random effect. Model selection was performed by stepwise removal of nonsignificant factors by comparing models of decreasing complexity based on Akaike information criterion, with the stipulation that body size was always included when eyespan remained in the model.

#### Fertility

To test if females mated to SR or ST males differed in fertility, we modelled total and proportion fertility as functions of mating group (one female or five females), body size, eyespan, genotype and the interaction between mating group and genotype, in generalised linear mixed effects models (GLMMs) including batch as a random effect.

To determine which reproductive traits were important predictors of fertility, total and proportion fertility were modelled as functions of mating group, body size, eyespan, testis size, accessory gland size and genotype, and up to all three-way interactions in

GLMMs including batch as a random effect. Model selection was performed based on Akaike information criterion as above, with the stipulation that body size was always included when eyespan, testis size or accessory gland size remained in the model.

Total fertility was modelled in a GLMM using a Poisson error distribution and a log link function, while proportion fertility was modelled using a binomial error distribution and logit link function. These data were over dispersed, and an observation-level random effect was added to account for this (Harrison 2014). Egg count data were only used if males had a minimum of 11 days of egg collections. Data were also excluded where females laid no eggs or fertility was <2%.

##### *Mating frequency*

We tested if there was a difference in the likelihood of SR and ST males mating at least once (coded as 1 or 0). This response was analysed as a function of genotype in a GLMM with a binomial distribution and logit link function, and batch as a random effect. The total number of copulations of those males that mated at least once was modelled as a function of male genotype in a GLMM using a Poisson error distribution and a log link function, with batch as a random effect.

### 6 Model tables and effect sizes

#### 6.1 SR trait size

##### Body size (thorax)

```r
lm(thorax^2 ~ genotype)
```

| | Sum Sq | Df | F value | Pr(>F) |
|---|---|---|---|---|
| genotype | 3.686 | 1 | 8.745 | 0.003 |
| Residuals | 150.490 | 357 | | |

| | Estimate | Std. Error | t value |
|---|---|---|---|
| (Intercept) | 5.479 | 0.042 | 130.179 |
| genotypeSR | -0.214 | 0.072 | -2.957 |
| N = 359 | | | |

##### Eyespan

```r
lm(eyespan ~ thorax + genotype)
```

| | Sum Sq | Df | F value | Pr(>F) |
|---|---|---|---|---|
| (Intercept) | 9.396 | 1 | 77.309 | <0.001 |
| thorax | 22.642 | 1 | 186.296 | <0.001 |
| genotype | 0.713 | 1 | 5.868 | 0.016 |

| | Sum Sq | Df | F value | Pr(>F) |
|---|---|---|---|---|
| thorax:genotype | 0.507 | 1 | 4.175 | 0.042 |
| Residuals | 43.145 | 355 | | |

| | Estimate | Std. Error | t value |
|---|---|---|---|
| (Intercept) | 3.295 | 0.375 | 8.793 |
| thorax | 2.186 | 0.160 | 13.649 |
| genotypeSR | -1.541 | 0.636 | -2.422 |
| thorax:genotypeSR | 0.563 | 0.275 | 2.043 |
| N = 359 | | | |

#### Eyespan (absolute)

```
lm(eyespan ~ genotype)
```

| | Sum Sq | Df | F value | Pr(>F) |
|---|---|---|---|---|
| genotype | 10.042 | 1 | 42.631 | <0.001 |
| Residuals | 84.090 | 357 | | |

| | Estimate | Std. Error | t value |
|---|---|---|---|
| (Intercept) | 8.402 | 0.031 | 267.072 |
| genotypeSR | -0.354 | 0.054 | -6.529 |
| N = 359 | | | |

#### Testes

```r
lmer(testis ~ females + thorax + residual_eyespan + genotype + (1 | batch))
```

| | F | Df | Df.res | Pr(>F) |
|---|---|---|---|---|
| females | 4.179 | 2 | 274.118 | 0.016 |
| thorax | 6.697 | 1 | 283.710 | 0.010 |
| residual_eyespan | 15.354 | 1 | 283.989 | <0.001 |
| genotype | 99.982 | 1 | 282.778 | <0.001 |

| | Estimate | Std. Error | t value |
|---|---|---|---|
| (Intercept) | 0.568 | 0.327 | 1.737 |
| females1 | 0.131 | 0.047 | 2.802 |
| females5 | 0.056 | 0.047 | 1.177 |
| thorax | 0.358 | 0.137 | 2.608 |
| residual_eyespan | 0.224 | 0.057 | 3.947 |
| genotypeSR | 0.409 | 0.041 | 10.052 |
| N = 290 | | | |

#### Testes (absolute)

```r
lmer(testis ~ females + genotype +  (1 | batch))
```

| | F | Df | Df.res | Pr(>F) |
|---|---|---|---|---|
| females | 3.937 | 2 | 274.529 | 0.021 |
| genotype | 73.796 | 1 | 280.158 | <0.001 |

| | Estimate | Std. Error | t value |
|---|---|---|---|
| (Intercept) | 1.426 | 0.053 | 26.792 |
| females1 | 0.130 | 0.048 | 2.703 |
| females5 | 0.053 | 0.048 | 1.093 |
| genotypeSR | 0.337 | 0.039 | 8.615 |
| N = 290 | | | |

#### Accessory glands

```
lmer(accessory_gland ~ thorax + residual_eyespan + genotype  +
    (1 | batch))
```

| | F | Df | Df.res | Pr(>F) |
|---|---|---|---|---|
| thorax | 0.693 | 1 | 335.628 | 0.406 |
| residual_eyespan | 8.971 | 1 | 335.691 | 0.003 |
| genotype | 7.801 | 1 | 334.029 | 0.006 |

| | Estimate | Std. Error | t value |
|---|---|---|---|
| (Intercept) | 0.261 | 0.101 | 2.590 |

| | Estimate | Std. Error | t value |
|---|---|---|---|
| thorax | 0.036 | 0.043 | 0.836 |
| residual_eyespan | 0.052 | 0.017 | 3.016 |
| genotypeSR | -0.036 | 0.013 | -2.804 |
| N = 340 | | | |

#### Accessory glands (absolute)

```
lmer(accessory_gland ~ genotype  + (1 | batch))
```

| | F | Df | Df.res | Pr(>F) |
|---|---|---|---|---|
| genotype | 16.353 | 1 | 335.356 | <0.001 |

| | Estimate | Std. Error | t value |
|---|---|---|---|
| (Intercept) | 0.349 | 0.011 | 31.371 |
| genotypeSR | -0.049 | 0.012 | -4.059 |
| N = 340 | | | |

#### Eyespan (residual) and accessory glands (residual) with testes

For these analyses, we are interested in the relationship between residual eyespan (fig. A1) or accessory gland size (fig. A2), and residual testis size within each genotype. Residual values are calculated for each genotype separately (using a linear model regressing thorax size on each of these traits).

```
lm(residual_eyespan_geno ~ residual_testis_geno * genotype)
```

| | Sum Sq | Df | F value | Pr(>F) |
|---|---|---|---|---|
| residual_testis_geno | 2.023 | 1 | 19.892 | <0.001 |
| genotype | 0.035 | 1 | 0.341 | 0.560 |
| residual_testis_geno:genotype | 0.535 | 1 | 5.261 | 0.023 |
| Residuals | 29.092 | 286 | | |

| | Estimate | Std. Error | t value |
|---|---|---|---|
| (Intercept) | -0.021 | 0.023 | -0.877 |
| residual_testis_geno | 0.388 | 0.082 | 4.739 |
| genotypeSR | 0.023 | 0.039 | 0.584 |
| residual_testis_geno:genotypeSR | -0.260 | 0.113 | -2.294 |
| N = 290 | | | |

```
lmer(residual_accessory_gland_geno ~ residual_testis_geno * genotype
    + (1 | batch))
```

| | F | Df | Df.res | Pr(>F) |
|---|---|---|---|---|
| (Intercept) | 0.171 | 1 | 18.781 | 0.684 |
| residual_testis_geno | 26.008 | 1 | 274.418 | <0.001 |
| genotype | 0.983 | 1 | 272.229 | 0.322 |
| residual_testis_geno:genotype | 8.375 | 1 | 268.066 | 0.004 |

| | Estimate | Std. Error | t value |
|---|---|---|---|
| (Intercept) | -0.005 | 0.013 | -0.413 |
| residual_testis_geno | 0.135 | 0.026 | 5.122 |
| genotypeSR | 0.012 | 0.012 | 0.995 |
| residual_testis_geno:genotypeSR | -0.102 | 0.035 | -2.899 |
| N = 281 | | | |

#### 6.2 SR fertility

##### SR total fertility

```
glmer(fertile_eggs ~ females + thorax + residual_eyespan +  genotype +
    females * genotype + (1 | batch) + (1 | OLRE))
```

| | Chisq | Df | Pr(>Chisq) |
|---|---|---|---|
| females | 43.698 | 1 | <0.001 |
| thorax | 0.688 | 1 | 0.407 |
| residual_eyespan | 1.439 | 1 | 0.230 |
| genotype | 2.416 | 1 | 0.120 |
| females:genotype | 0.591 | 1 | 0.442 |

| | Estimate | Std. Error | z value |
|---|---|---|---|
| (Intercept) | 3.893 | 0.106 | 36.880 |
| females5 | 0.853 | 0.144 | 5.928 |

| | Estimate | Std. Error | z value |
|---|---|---|---|
| thorax | 0.051 | 0.061 | 0.830 |
| residual_eyespan | 0.076 | 0.063 | 1.199 |
| genotypeSR | 0.316 | 0.190 | 1.666 |
| females5:genotypeSR | -0.201 | 0.262 | -0.769 |
| N = 215 | | | |

#### SR total fertility (absolute)

```r
glmer(fertile_eggs ~ females * genotype + (1 | batch) + (1 | OLRE))
```

| | Chisq | Df | Pr(>Chisq) |
|---|---|---|---|
| females | 43.740 | 1 | <0.001 |
| genotype | 0.926 | 1 | 0.336 |

| | Estimate | Std. Error | z value |
|---|---|---|---|
| (Intercept) | 3.962 | 0.101 | 39.140 |
| females5 | 0.786 | 0.119 | 6.614 |
| genotypeSR | 0.128 | 0.133 | 0.963 |
| N = 215 | | | |

#### SR proportion fertility

```
glmer(cbind(fert, unfert) ~ females + thorax + residual_eyespan +
    genotype + females * genotype + (1|batch) + (1|OLRE))
```

| | Chisq | Df | Pr(>Chisq) |
|---|---|---|---|
| females | 6.021 | 1 | 0.014 |
| thorax | 1.268 | 1 | 0.260 |
| residual_eyespan | 0.017 | 1 | 0.895 |
| genotype | 2.469 | 1 | 0.116 |
| females:genotype | 1.377 | 1 | 0.241 |

| | Estimate | Std. Error | z value |
|---|---|---|---|
| (Intercept) | 1.759 | 0.195 | 9.006 |
| females5 | -0.345 | 0.249 | -1.388 |
| thorax | 0.130 | 0.115 | 1.126 |
| residual_eyespan | 0.015 | 0.112 | 0.132 |
| genotypeSR | 0.673 | 0.345 | 1.950 |
| females5:genotypeSR | -0.532 | 0.453 | -1.173 |
| N = 215 | | | |

#### SR proportion fertility (absolute)

```
glmer(cbind(fert, unfert) ~ females * genotype + (1|batch) + (1|OLRE))
```

| | Chisq | Df | Pr(>Chisq) |
|---|---|---|---|
| females | 6.910 | 1 | 0.009 |
| genotype | 2.156 | 1 | 0.142 |
| females:genotype | 1.036 | 1 | 0.309 |

| | Estimate | Std. Error | z value |
|---|---|---|---|
| (Intercept) | 1.812 | 0.193 | 9.376 |
| females5 | -0.402 | 0.243 | -1.655 |
| genotypeSR | 0.563 | 0.319 | 1.762 |
| females5:genotypeSR | -0.456 | 0.448 | -1.018 |
| N = 215 | | | |

#### SR fecundity

```
glmer(total.eggs ~ females + thorax + residual_eyespan + genotype +
    females * genotype + (1 |OLRE) + (1 | batch))
```

| | Chisq | Df | Pr(>Chisq) |
|---|---|---|---|
| females | 78.719 | 1 | <0.001 |
| thorax | 0.297 | 1 | 0.586 |

| | Chisq | Df | Pr(>Chisq) |
|---|---|---|---|
| residual_eyespan | 1.562 | 1 | 0.211 |
| genotype | 1.079 | 1 | 0.299 |
| females:genotype | 0.096 | 1 | 0.757 |

| | Estimate | Std. Error | z value |
|---|---|---|---|
| (Intercept) | 4.303 | 0.075 | 57.269 |
| females5 | 0.776 | 0.103 | 7.562 |
| thorax | 0.024 | 0.044 | 0.545 |
| residual_eyespan | 0.057 | 0.045 | 1.250 |
| genotypeSR | 0.132 | 0.136 | 0.975 |
| females5:genotypeSR | -0.058 | 0.187 | -0.310 |
| N = 215 | | | |

#### Organs total fertility

```
glmer(fertile_eggs ~ females + thorax + residual_eyespan +
    residual_testis + residual_accessory_gland +
    residual_eyespan:residual_accessory_gland + (1 | batch) +
    (1 | OLRE))
```

| | Chisq | Df | Pr(>Chisq) |
|---|---|---|---|
| (Intercept) | 2188.421 | 1 | <0.001 |
| females | 43.547 | 1 | <0.001 |

| | Chisq | Df | Pr(>Chisq) |
|---|---|---|---|
| thorax | 0.509 | 1 | 0.475 |
| residual_eyespan | 0.208 | 1 | 0.648 |
| residual_testis | 6.216 | 1 | 0.013 |
| residual_accessory_gland | 0.032 | 1 | 0.858 |
| residual_eyespan:residual_accessory_gland | 7.133 | 1 | 0.008 |

| | Estimate | Std. Error | z value |
|---|---|---|---|
| (Intercept) | 4.096 | 0.088 | 46.781 |
| females5 | 0.762 | 0.115 | 6.599 |
| thorax | 0.046 | 0.064 | 0.714 |
| residual_eyespan | 0.029 | 0.063 | 0.456 |
| residual_testis | 0.148 | 0.059 | 2.493 |
| residual_accessory_gland | 0.011 | 0.060 | 0.179 |
| residual_eyespan:residual_accessory_gland | 0.165 | 0.062 | 2.671 |
| N = 165 | | | |

#### Organs total fertility: testis and genotype interaction

```
glmer(fertile_eggs ~ females + thorax + residual_eyespan +
    residual_testis * genotype + (1 | batch) + (1 | OLRE))
```

| | Chisq | Df | Pr(>Chisq) |
|---|---|---|---|
| (Intercept) | 1483.845 | 1 | <0.001 |

| | Chisq | Df | Pr(>Chisq) |
|---|---|---|---|
| females | 43.563 | 1 | <0.001 |
| thorax | 0.068 | 1 | 0.794 |
| residual_eyespan | 0.008 | 1 | 0.927 |
| residual_testis | 4.159 | 1 | 0.041 |
| genotype | 0.018 | 1 | 0.895 |
| residual_testis:genotype | 0.164 | 1 | 0.686 |

| | Estimate | Std. Error | z value |
|---|---|---|---|
| (Intercept) | 4.119 | 0.107 | 38.521 |
| females5 | 0.778 | 0.118 | 6.600 |
| thorax | 0.017 | 0.065 | 0.261 |
| residual_eyespan | -0.007 | 0.075 | -0.091 |
| residual_testis | 0.203 | 0.099 | 2.039 |
| genotypeSR | -0.023 | 0.175 | -0.132 |
| residual_testis:genotypeSR | -0.056 | 0.138 | -0.404 |
| N = 173 | | | |

#### Organs total fertility (absolute)

```
glmer(fertile_eggs ~ females + testis + accessory_gland +
    (1 | batch) + (1 | OLRE))
```

| | Chisq | Df | Pr(>Chisq) |
|---|---|---|---|
| (Intercept) | 2260.899 | 1 | <0.001 |
| females | 40.426 | 1 | <0.001 |
| testis | 5.897 | 1 | 0.015 |
| accessory_gland | 0.260 | 1 | 0.610 |

| | Estimate | Std. Error | z value |
|---|---|---|---|
| (Intercept) | 4.132 | 0.087 | 47.549 |
| females5 | 0.747 | 0.118 | 6.358 |
| testis | 0.148 | 0.061 | 2.428 |
| accessory_gland | 0.031 | 0.060 | 0.510 |
| N = 165 | | | |

#### Organs proportion fertility

```
glmer(cbind(fert, unfert) ~ females + thorax + residual_eyespan +
    residual_testis + residual_eyespan:residual_testis +  (1|batch) +
    (1|OLRE))
```

| | Chisq | Df | Pr(>Chisq) |
|---|---|---|---|
| (Intercept) | 205.835 | 1 | <0.001 |
| females | 9.620 | 1 | 0.002 |
| thorax | 1.816 | 1 | 0.178 |
| residual_eyespan | 0.724 | 1 | 0.395 |

| | Chisq | Df | Pr(>Chisq) |
|---|---|---|---|
| residual_testis | 16.646 | 1 | <0.001 |
| residual_accessory_gland | 0.160 | 1 | 0.689 |
| residual_eyespan:residual_testis | 3.867 | 1 | 0.049 |

| | Estimate | Std. Error | z value |
|---|---|---|---|
| (Intercept) | 2.156 | 0.150 | 14.347 |
| females5 | -0.601 | 0.194 | -3.102 |
| thorax | 0.138 | 0.102 | 1.348 |
| residual_eyespan | -0.093 | 0.109 | -0.851 |
| residual_testis | 0.421 | 0.103 | 4.080 |
| residual_accessory_gland | 0.041 | 0.103 | 0.400 |
| residual_eyespan:residual_testis | 0.222 | 0.113 | 1.966 |
| N = 165 | | | |

#### Organs proportion fertility: testis and genotype interaction

```
glmer(cbind(fert, unfert) ~ females + thorax + residual_eyespan +
    residual_testis * genotype +  (1|batch) + (1|OLRE))
```

| | Chisq | Df | Pr(>Chisq) |
|---|---|---|---|
| (Intercept) | 133.483 | 1 | <0.001 |
| females | 6.268 | 1 | 0.012 |
| thorax | 2.421 | 1 | 0.120 |

| | Chisq | Df | Pr(>Chisq) |
|---|---|---|---|
| residual_eyespan | 2.674 | 1 | 0.102 |
| residual_testis | 12.198 | 1 | <0.001 |
| genotype | 0.260 | 1 | 0.610 |
| residual_eyespan:residual_testis | 4.889 | 1 | 0.027 |
| residual_testis:genotype | 0.617 | 1 | 0.432 |

| | Estimate | Std. Error | z value |
|---|---|---|---|
| (Intercept) | 2.141 | 0.185 | 11.554 |
| females5 | -0.497 | 0.199 | -2.504 |
| thorax | 0.162 | 0.104 | 1.556 |
| residual_eyespan | -0.200 | 0.122 | -1.635 |
| residual_testis | 0.612 | 0.175 | 3.493 |
| genotypeSR | -0.148 | 0.290 | -0.510 |
| residual_eyespan:residual_testis | 0.257 | 0.116 | 2.211 |
| residual_testis:genotypeSR | -0.187 | 0.238 | -0.786 |
| N = 173 | | | |

#### Organs proportion fertility (absolute)

```
glmer(cbind(fert, unfert) ~ females + testis + accessory_gland +
    (1|batch) + (1|OLRE))
```

| | Chisq | Df | Pr(>Chisq) |
|---|---|---|---|
| (Intercept) | 210.014 | 1 | <0.001 |
| females | 10.084 | 1 | 0.001 |
| testis | 18.837 | 1 | <0.001 |
| accessory_gland | 0.140 | 1 | 0.708 |

| | Estimate | Std. Error | z value |
|---|---|---|---|
| (Intercept) | 2.191 | 0.151 | 14.492 |
| females5 | -0.619 | 0.195 | -3.176 |
| testis | 0.452 | 0.104 | 4.340 |
| accessory_gland | 0.039 | 0.105 | 0.374 |
| N = 165 | | | |

#### Organs fecundity

```
glmer(total.eggs ~ females + thorax + residual_eyespan +
    residual_accessory_gland + genotype +
    residual_eyespan:residual_accessory_gland + (1 |OLRE) + (1 | batch))
```

| | Chisq | Df | Pr(>Chisq) |
|---|---|---|---|
| (Intercept) | 3926.294 | 1 | <0.001 |
| females | 70.210 | 1 | <0.001 |
| thorax | 0.130 | 1 | 0.719 |
| residual_eyespan | 0.007 | 1 | 0.934 |

| | Chisq | Df | Pr(>Chisq) |
|---|---|---|---|
| residual_accessory_gland | 0.048 | 1 | 0.826 |
| residual_eyespan:residual_accessory_gland | 3.995 | 1 | 0.046 |

| | Estimate | Std. Error | z value |
|---|---|---|---|
| (Intercept) | 4.369 | 0.070 | 62.660 |
| females5 | 0.717 | 0.086 | 8.379 |
| thorax | -0.018 | 0.049 | -0.360 |
| residual_eyespan | -0.004 | 0.048 | -0.082 |
| residual_accessory_gland | 0.010 | 0.045 | 0.219 |
| residual_eyespan:residual_accessory_gland | 0.081 | 0.041 | 1.999 |
| N = 199 | | | |

#### 6.3 SR mating frequency

##### Mated at least once

```
glmer(mated ~ genotype + (1|batch), family = binomial)
```

| | Chisq | Df | Pr(>Chisq) |
|---|---|---|---|
| genotype | 1.665 | 1 | 0.197 |

| | Estimate | Std. Error | z value |
|---|---|---|---|
| (Intercept) | 1.830 | 0.336 | 5.44 |
| genotypeSR | -0.501 | 0.389 | -1.29 |
| N = 193 | | | |

#### Total copulations

```
glmer(total_matings ~ genotype + (1|batch), family = poisson)
```

| | Chisq | Df | Pr(>Chisq) |
|---|---|---|---|
| genotype | 6.304 | 1 | 0.012 |

| | Estimate | Std. Error | z value |
|---|---|---|---|
| (Intercept) | 1.246 | 0.072 | 17.342 |
| genotypeSR | -0.235 | 0.093 | -2.511 |
| N = 157 | | | |

### 7 Figures

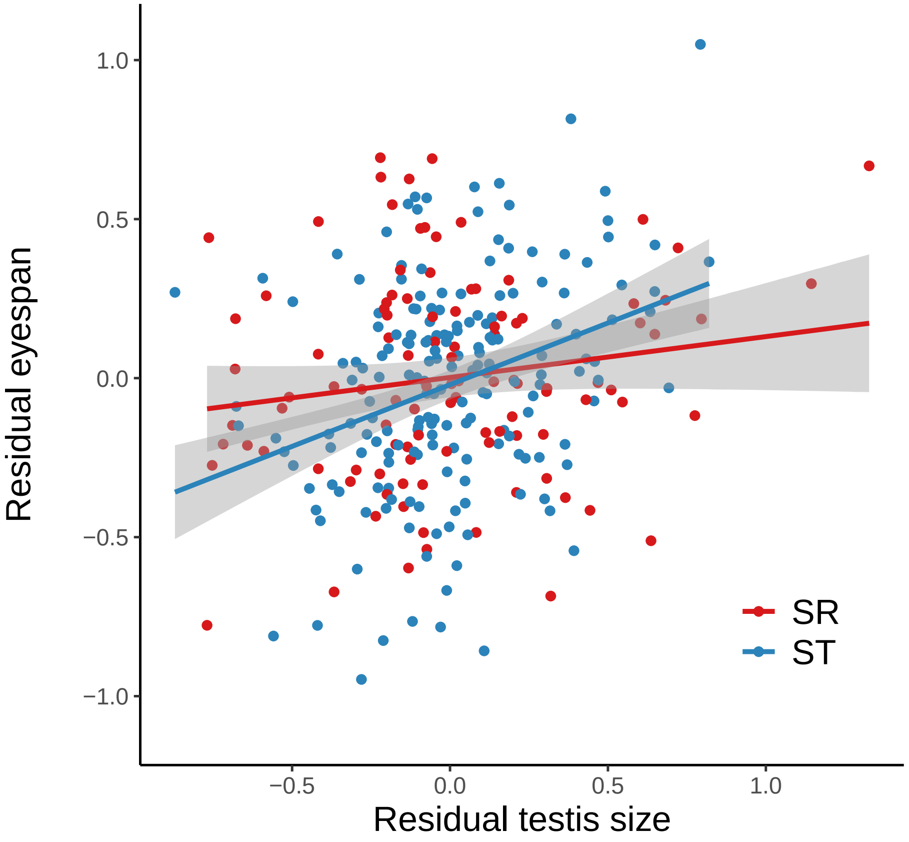

**Figure A1:** Male residual eyespan for SR (red) and ST (blue) males, regressed on male residual testis size. ST males with large residual testis also have large residual eyespan, but this is less so for SR males (interaction P = 0.023). Residuals were calculated separately for each genotype by regressing thorax size on eyespan or testis size. Grey shading shows ± s.e.

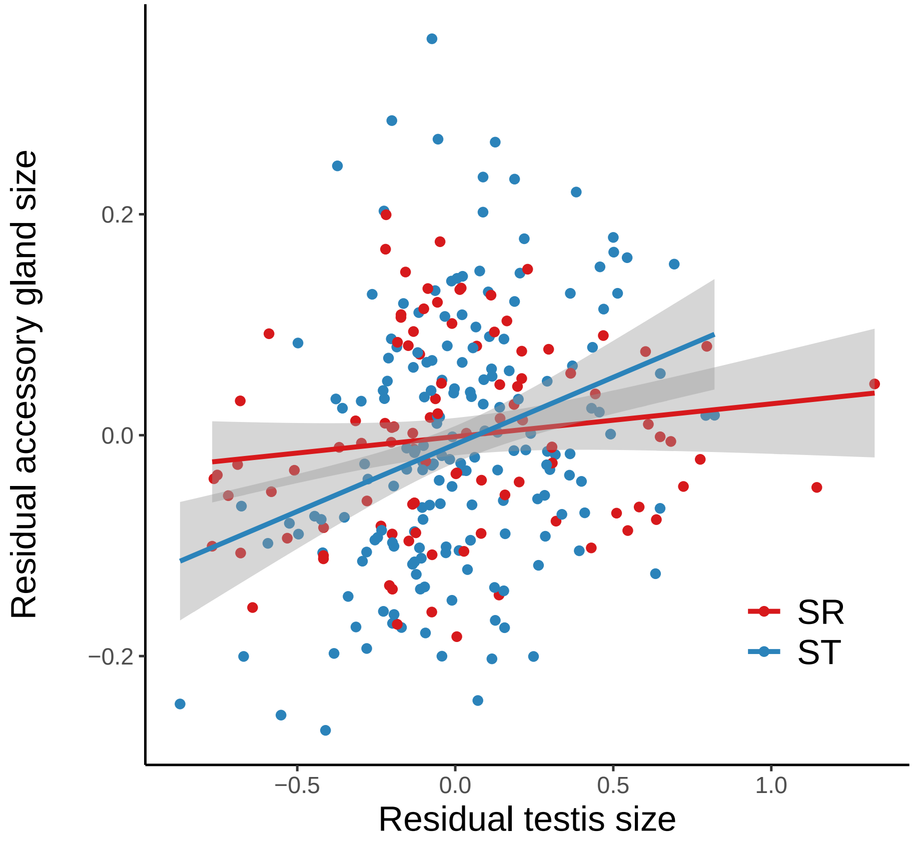

**Figure A2:** Male residual accessory gland size for SR (red) and ST (blue) males, regressed on male residual testis size. ST males with large residual testis also have large residual accessory gland size, but this is less so for SR males (interaction P = 0.004). Residuals were calculated separately for each genotype by regressing thorax size on accessory gland size or testis size. Grey shading shows ± s.e.
